# An Alignment-free Method for Phylogeny Estimation using Maximum Likelihood

**DOI:** 10.1101/2019.12.13.875526

**Authors:** Tasfia Zahin, Md. Hasin Abrar, Mizanur Rahman, Tahrina Tasnim, Md. Shamsuzzoha Bayzid, Atif Rahman

## Abstract

While alignment has traditionally been the primary approach for establishing homology prior to phylogenetic inference, alignment-free methods offer a simplified alternative, particularly beneficial when handling genome-wide data involving long sequences and complex events such as rearrangements. Moreover, alignment-free methods become crucial for data types like genome skims, where assembly is impractical. However, despite these benefits, alignment-free techniques have not gained widespread acceptance since they lack the accuracy of alignment-based techniques, primarily due to their reliance on simplified models of pairwise distance calculation. Here, we present a likelihood based alignment-free technique for phylogenetic tree construction. We encode the presence or absence of *k*-mers in genome sequences in a binary matrix, and estimate phylogenetic trees using a maximum likelihood approach. We analyze the performance of our method on seven real datasets and compare the results with the state of the art alignment-free methods. Results suggest that our method is competitive with existing alignment-free tools. This indicates that maximum likelihood based alignment-free methods may in the future be refined to outperform alignment-free methods relying on distance calculation as has been the case in the alignment-based setting. A likelihood based alignment-free method for phylogeny estimation is implemented for the first time in a software named Peafowl, which is available at: https://github.com/hasin-abrar/Peafowlrepo.

## 1. Introduction

A phylogenetic tree depicts the evolutionary history of a given set of species. Efficient and accurate construction of phylogenies from genome data is one of the most important problems in biology and is a major research focus in bioinformatics and systematics. Phylogeny construction methods can be broadly classified into two groups: distance based and character based. Distance based methods compute the distances from the genomic sequences of each pair of species to construct a distance matrix. Tree construction algorithms are then applied to this matrix to estimate the tree topology. Popular distance based methods include *UPGMA* [1], *neighbor-joining* [2], etc. They are fast and can handle many sequences but their performance is dependent on the accuracy of the distance matrix. Character based methods, on the other hand, make use of the sequences typically in the form of a multiple sequence alignment (MSA). *Maximum parsimony* [3] is a character based approach where a character matrix is taken as input and the best tree under the maximum parsimony criterion is the one that minimizes the number of changes in the nucleotide sequences over time. *Maximum likelihood* [4], a probabilistic character based approach, uses specific models of sequence evolution to find a tree that maximizes the likelihood of observing the set of input sequences. This approach is quite realistic in nature and suitable for species that vary widely in terms of similarity unlike the maximum parsimony approach.

Previous studies indicate that, in general, maximum likelihood approaches are superior in terms of performance over distance based methods. Maximum likelihood based methods were observed to estimate correct trees better than the neighbor joining method when the underlying assumptions behind the methods are not satisfied [5]. In addition, maximum likelihood based methods are also more robust than distance based methods using least square criterion [6].

However, in the alignment-based paradigm, both distance based and character based approaches require prior alignment of input sequences. The quality of alignment greatly affects the resulting phylogeny. Sequence alignment is memory and time consuming, and hence is difficult to scale to large sequences and whole genomes. Moreover, finding an optimal multiple sequence alignment is known to be computationally intractable as the number of possible alignments increases exponentially with increasing sequence lengths [7]. Furthermore, alignment-based methods assume a preserved linear order of homology, and therefore the presence of rearrangement events, such as translocation, inversion, etc. within whole genome sequences complicates sequence alignment – making it even more challenging to construct accurate phylogenetic trees from whole genomes [7].

To overcome the aforementioned difficulties, phylogenetic analyses that are not confined to alignment needs are gaining increasing attention, saving substantial time and memory in the phylogeny estimation process. The methods are collectively known as *alignment-free* methods. They are robust to rearrangement events and suitable for phylogeny estimation from large sequences and even whole genomes. However, despite their practical advantages, alignment-free techniques have not demonstrated the same level of accuracy as alignment-based methods. It is important to acknowledge that we do not anticipate alignment-free methods to match the accuracy of alignment-based methods, particularly when dealing with small, rearrangement-free sequences such as single genes. This is because alignment-free methods still require effective strategies for handling homology, a challenge that is no less complex than alignment itself.

A multitude of recently developed alignment-free methods have been comprehensively reviewed in [8, 9]. Among these, *co-phylog* [10] searches for short alignments of fixed length in the sequences allowing a mismatch in the middle. Evolutionary distances are calculated from these sub-sequences, followed by tree generation. *andi* [11] looks for mismatches surrounded by long exact matches. Counts of mismatches are used to estimate the number of substitutions between two sequences. *mash* [12] is based on the MinHash technique to find representative sketches of sequences from which Jaccard indices are estimated as a distance measure. *Multi-SpaM* [13] uses the *Space Word Match (SWM)* [14] approach to identify quartet groups, *i*.*e*. a group of four space words with matching nucleotides at the match positions and probable mismatches at the don’t care positions.

However, despite their potential, alignment-free methods have not yet been found to be as accurate as alignment-based methods. Majority of the alignment-free methods developed so far are distance based and hence do not allow model based phylogeny estimation that are known to be more robust than the former. Höhl and Ragan [15] proposed a Bayesian approach for phylogeny inference based on the existence of *k*-mers (contiguous subsequences of length *k*) in the sequences.

In this paper, motivated by the observation that methods using maximum likelihood outperform distance based methods in the alignment-based setting, we present an alignment-free method for phylogenetic tree construction that utilizes maximum likelihood estimation. We first construct a matrix encoding the presence or absence of *k*-mers within the sequences, and then use an existing model for binary traits to construct a phylogeny that maximizes like-lihood. The method is implemented in a tool called Peafowl (**P**hylogeny **E**stimation through **A**lignment **F**ree **O**ptimization **W**ith **L**ikelihood). We analyze the performance of our method by applying it on seven real datasets, including datasets from the AFproject [9] which is widely used for assessing alignment-free tools.

## 2. Methods

An overview of phylogenetic tree estimation using Peafowl is shown in Figure 1. The method consists of four major steps. First, the set of *k*-mers present in each input sequence is generated. Second, a binary matrix is constructed, which encapsulates the presence or absence of the *k*-mers within the sequences. Third, a suitable value of *k* is chosen based on entropy values. Finally, a phylogenetic tree is constructed using maximum likelihood estimation. A sketch of the steps is presented in Algorithm 1 and described in more detail in the following sections.

**Figure 1:**
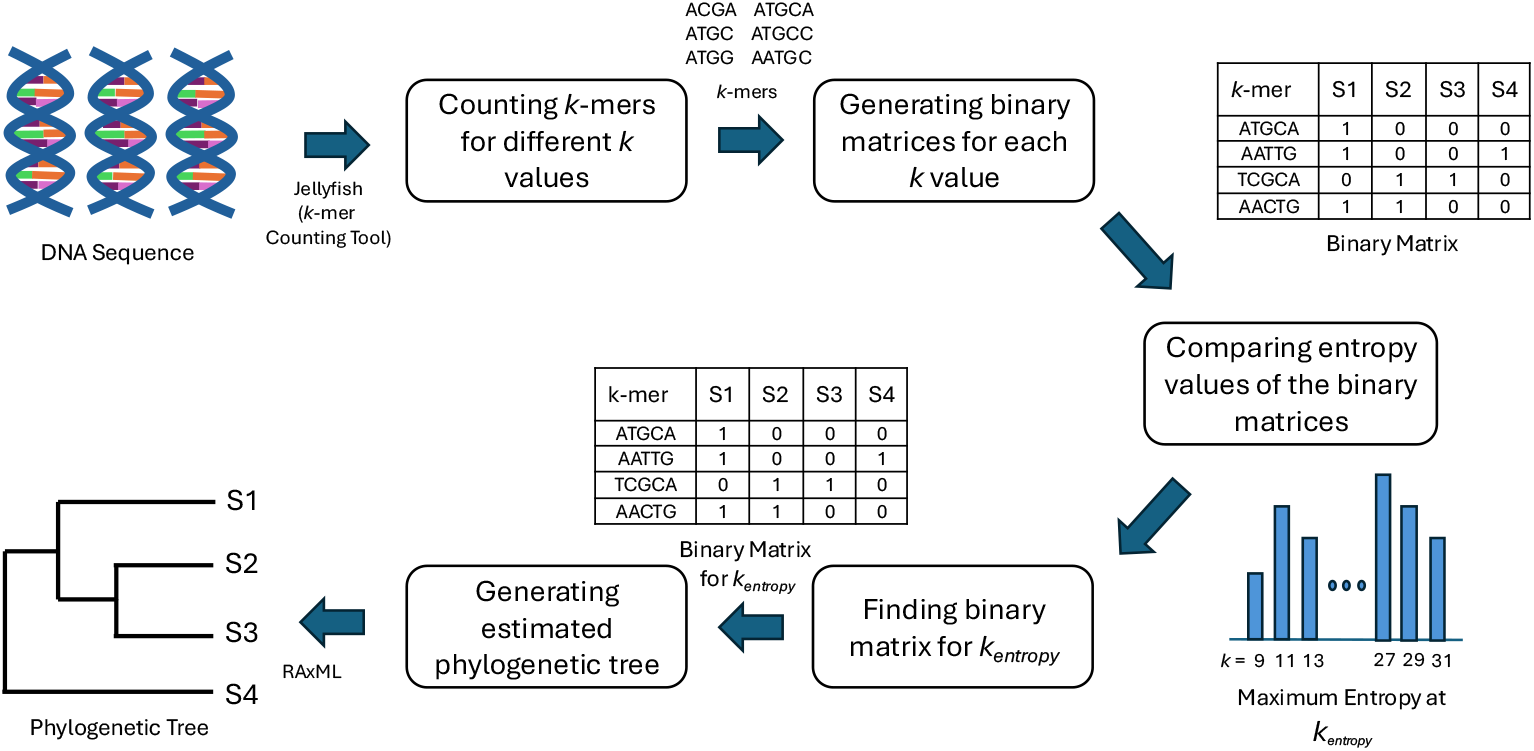
Overview of phylogenetic tree estimation using Peafowl. At the beginning, *k*-mers of various sizes are listed from the input sequences. Then separate binary matrices are produced using these *k*-mers. From the binary matrices of different *k*-mer sizes, an appropriate *k*-mer length (*k*_*entropy*_) is chosen based on cumulative entropy values. Lastly, the binary matrix corresponding to *k*_*entropy*_ is provided as input to RAxML for the estimation of the phylogenetic tree.

**Figure 2:**
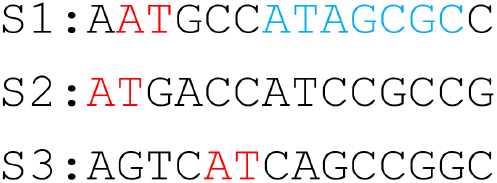
Choosing *k*-mer lengths. Existence of *k*-mers depends on the length. In this figure, the *k*-mer AT of length 2 is found in all 3 taxa. However, the *k*-mer ATAGCGC of length 7 is found only in the source taxon (T1).

### 2.1. Generating k-mers

The first step in Peafowl is to generate the lists of *k*-mers present in the input sequences. *k*-mers are generated from the input DNA sequences using Jellyfish [16] for odd values of *k* ranging from 9 to 31 (more details in Subsection Finding an appropriate *k*-mer length). As DNA is double-stranded and the sequences in the two strands are complements of each other, the input sequences can be from one strand or from both. In the latter case, it is more appropriate to consider a *k*-mer and its reverse complement as the same during counting, usually referred to as canonical counting. In the former case, a *k*-mer and its reverse complement can be treated independently, commonly known as non-canonical counting. Our method is designed to work in both possible modes, allowing the user to choose how reverse complements should be treated during the *k*-mer counting step. However, all the results shown in this paper except one (see Subsection Horizontal gene transfer (HGT)) are obtained using the canonical counting mode as the assembled sequences may correspond to either strand of DNA.

### 2.2. Generating binary matrices

The next step is to construct a binary matrix denoting whether the generated *k*-mers are present in the given sequences or not. This matrix consists of only 0’s and 1’s. Its rows and columns represent the *k*-mers and the input species, respectively. An entry in the matrix contains 1 if the *k*-mer representing the row (or its reverse complement) exists in the sequence of the species representing the column and 0 otherwise. One such matrix is produced for each value of *k*. We use hashing for this particular task. *k*-mers are read from a file, and a unique index is generated for each of them. The *k*-mers are inserted into a hash table along with the identification numbers of species they come from. The hash table indices are accessed one after another while placing an appropriate value in the desired position of the matrix.

#### Algorithm 1

Phylogeny estimation using Peafowl

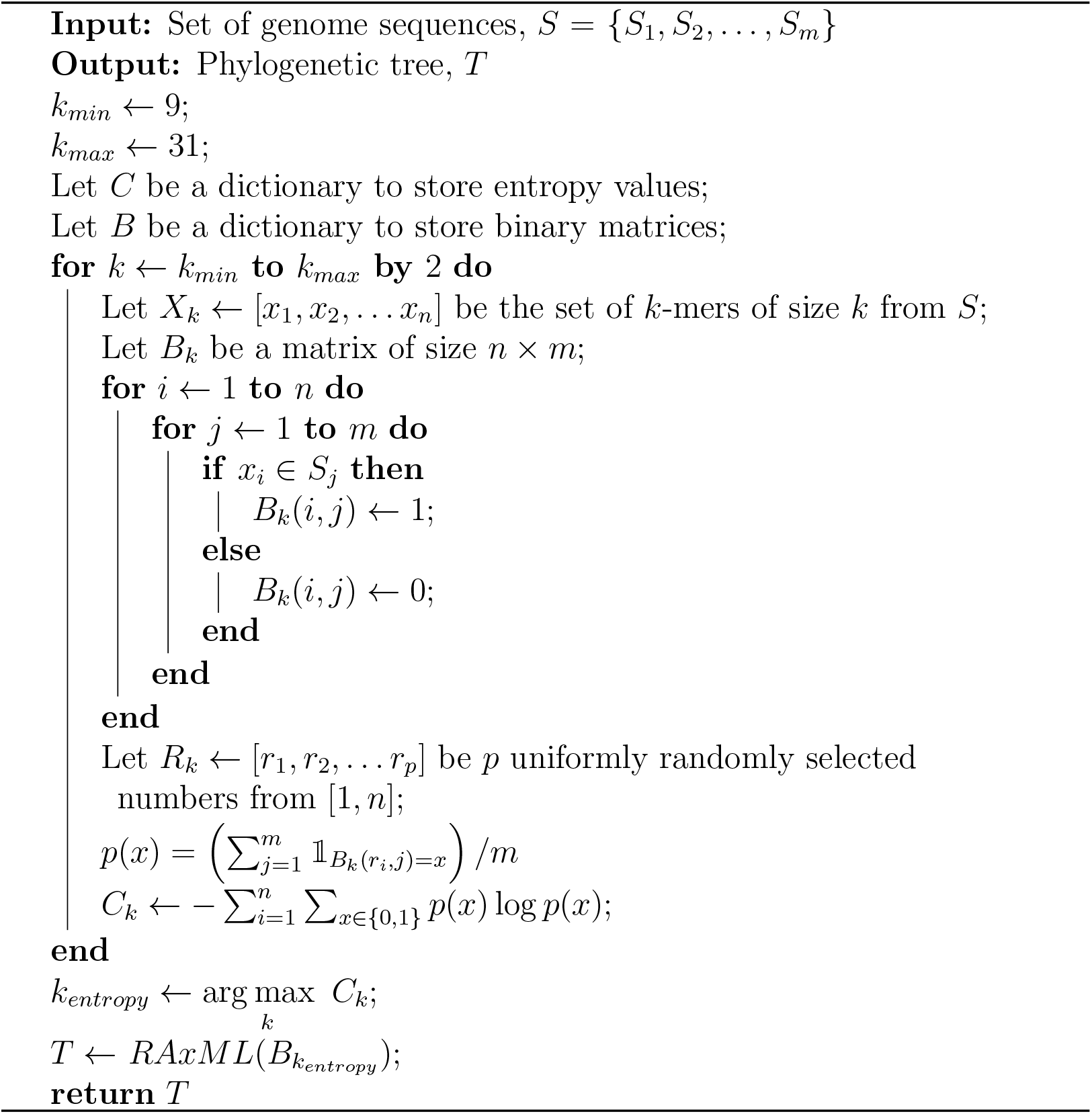

### 2.3. Finding an appropriate k-mer length

A number of approaches have been proposed by researchers to choose a proper *k*-mer length for alignment-free analysis. [17] applies a logarithmic function on input sequence lengths to calculate a suitable value of *k*. The limitation of this selection process is that it does not take into account how closely related the species are. *Slope-SpaM* [18] analyzes match probability to calculate lower and upper bounds on *k*-mer length. However, it does not serve the need for a specific value of *k* that our model requires. More specifically, our target is to find a value of *k* that would capture the most informative binary matrix for tree generation.

The genetic diversity among the different genome sequences can be modeled by the concept of entropy [19]. This concept has been previously used by several other sequence analysis approaches [20, 21]. We utilize this in our method for *k*-mer length selection. *k*-mers that can be found in almost all the species introduce many 1s in the matrix, while rare ones introduce many 0s. *k*-mers that do not fall in either of these extremities provide comparatively more information. A binary matrix rich in these types of *k*-mers will capture the relationship between species better than the others. Since entropy can capture the randomness in a system, we use this metric to compare the information content of the binary matrices, and choose an appropriate *k*-mer length. Cumulative entropy for any binary matrix is calculated using the following equation.

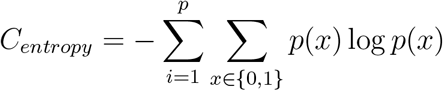

where

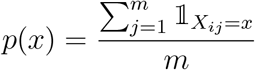

Here, *p* represents the number of *k*-mers used for entropy calculation, and *m* represents the total number of species. *X*_*ij*_ represents the state of an entry in the matrix corresponding to the *i*^*th*^ row and the *j*^*th*^ species, and can take values of either 0 or 1. Again, 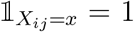 if *X*_*ij*_ = *x*, and 0 otherwise. The equation adds the entropy values of *p* randomly selected rows to get the cumulative entropy (*C*_*entropy*_) for any binary matrix. Here, *p* is empirically chosen to be 5000.

Empirical evidence suggests that *k* values less than 9 cause *k*-mers to be excessively abundant, while those greater than 31 often lead to the presence of *k*-mers in only one or a few sequences [18]. For even values of *k*, a *k*-mer and its reverse complement may become the same causing inconsistency in the non-canonical counting mode [22]. To address these issues and to reduce the computational complexity, binary matrices are created for odd *k*-mer lengths ranging from 9 to 31. *C*_*entropy*_ values from different *k*-mer lengths are compared and the value of *k* resulting in the maximum entropy is selected to be the most suitable one. We refer to this length as *k*_*entropy*_.

### 2.4. Generating phylogenetic trees

Once we have obtained *k*_*entropy*_, the final step is to construct a tree from the binary matrix corresponding to this length. This is done by providing the concerned matrix as input into a widely used tool for maximum likelihood phylogeny estimation named RAxML [23]. We use an existing model of substitution for binary traits BINGAMMA for our method. It is defined for binary data and assumes a gamma prior on the site mutation rates. The model takes in binary sequences and outputs an estimated tree topology, assuming sites to be independent. However, in reality, one character substitution in a sequence affects a number of neighboring *k*-mers at that site. This is why we focus on tree topology for now and leave branch length estimation as future work.

### 2.5. Implementation

Peafowl is implemented using C++ and shell scripts. In addition, it uses Jellyfish 2.2.4 for *k*-mer counting. A rigorous comparison of *k*-mer counting methods is presented by Zhang et al. [24]. We choose Jellyfish [16] on the basis of this comparison. The tool is fast, supports dynamic memory and is preferable for large genome sequences.

Peafowl also uses RAxML 8.2.4 for phylogeny estimation. RAxML stands for **R**andomized **Ax**elerated **M**aximum **L**ikelihood [23]. It is a popular phylogenetic analysis software that can handle large datasets and is useful for maximum likelihood based phylogeny inference.

## 3. Results

### 3.1. Datasets and benchmarking

We assess the performance of our method using seven real datasets. First, we analyze a 7 Primates dataset [8] and a Drosophila dataset from [25]. The 7 Primates dataset contains full mitochondrial genome sequences of 7 primates, and the Drosophila dataset consists of real genome skims of 14 Drosophila species subsampled to 100 Mb. We selected these datasets as the reference trees for these species are well established.

Next, we analyzed datasets from the AFproject [9] that have been widely used for benchmarking alignment-free methods. We selected the five real datasets under the Genome-based Phylogeny and Horizontal Gene Transfer categories that had assembled genomes. They include assembled sequences of 29 *E*.*coli/Shigella* strains [10], assembled mitochondrial genomes of 25 fish species of the suborder *Labroidei* [26], full genome sequences of 14 plant species [27], full genome sequences of 27 *E*.*coli/Shigella* strains [28], and full genome sequences of 8 Yersinia strains [28].

Genome sequences, benchmark trees, and results of the last five datasets were obtained from the AFproject [9]. For the primates and Drosophila datasets, sequences and the benchmark tree are obtained from [8] and [25], respectively. The primary performance metric used throughout this paper is the *Robinson Foulds (RF)* [29] distance. It gives a measure of the distance between two trees by counting the number of dissimilar partitions. The distance is divided by the maximum possible RF value to obtain the normalized RF distance (nRF). The smaller this score, the more congruent the estimated and the reference trees.

Our method is run on these datasets with the *-r* parameter (canonical *k*-mer counting) i.e. reverse complements are considered the same *k*-mer. The tree corresponding to *k*_*entropy*_ is treated as the final tree. The nRF distance between this tree and the reference tree is compared to those achieved by state-of-the-art methods from the AFproject [9]. The bench-marked methods include *FFP* [30, 31], *co-phylog* [10], *mash* [12], *Skmer* [25] *FSWM/Read-SpaM* [32, 33] *andi* [11], *phylonium* [34], *Multi-SpaM* [13], and *CAFE-cvtree* [35]. It has been observed that no single method benchmarked by AFproject [9] achieves the best scores across all datasets. The aforementioned methods include the top performers.

Some of the benchmarked tools generate a distance matrix as output and not a phylogenetic tree. Therefore, for the 7 primates and Drosophila datasets, we apply neighbor-joining and UPGMA implementation of MEGA-X [36] on the distance matrix to get the estimated trees, and find the RF distances using PHYLIP [37]. nRF values of benchmarking tools reported here are from trees produced by neighbor-joining. Results from UPGMA are available in the supplementary materials. We limit the scope of our work to phylogenetic trees upto a maximum of 30 species.

### 3.2. Selection of k-mer lengths

We first explore how the estimated trees vary with different *k*-mer lengths. The variation of nRF and entropy with change in *k*-mer size for the 7 Primates and Drosophila datasets are illustrated in Figure 3. Similar plots for the remaining datasets are shown in Supplementary Figures (S1 Figure - S5 Figure). We observe that, for the 7 Primates dataset, the minimum nRF distance of 0 and the maximum entropy is obtained when *k* equals 9. In all cases, we find that the lowest nRF distances occur at the *k*-mer lengths with the highest entropy i.e. *k*_*entropy*_. We observe that in Figure 3(a) there is a drop in nRF at *k*=23. This might be because the dataset contains only seven species, so a change in a single branch leads to a substantial decrease in the nRF value. In the subsequent sections, we only report nRF distances corresponding to the tree obtained using *k*_*entropy*_.

**Figure 3:**
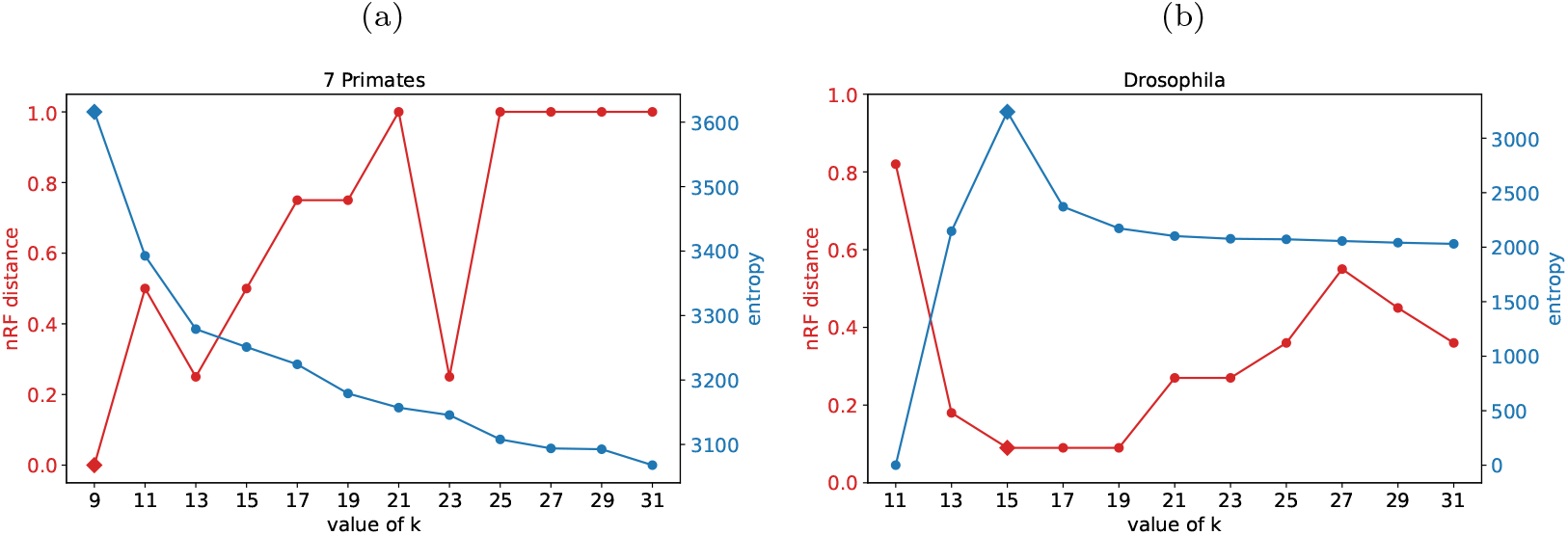
nRF and entropy vs. *k*-mer. Variation of normalized Robinson Foulds distance and entropy with change in *k*-mer length for (a) the 7-Primates dataset and (b) the Drosophila dataset. Diamond shaped markers represent values corresponding to *k*_*entropy*_.

### 3.3. 7 Primates and Drosophila Datasets

The nRF distances for Peafowl and other methods for the 7 Primates and Drosophila datasets are demonstrated in Figure 4. Peafowl along with a few other methods (e.g., andi, Multi-SpaM, FFP) correctly reconstructed the reference tree. It is worth noting that highly accurate methods like Mash, Skmer, and co-phylog placed Gibbon as the sister to hominine (gorillas, chim-panzees, and humans) and thus failed to reconstruct the well-established (orangutan, (gorilla, (chimpanzee, human))) relationship (see Fig. 5(a)).

**Figure 4:**
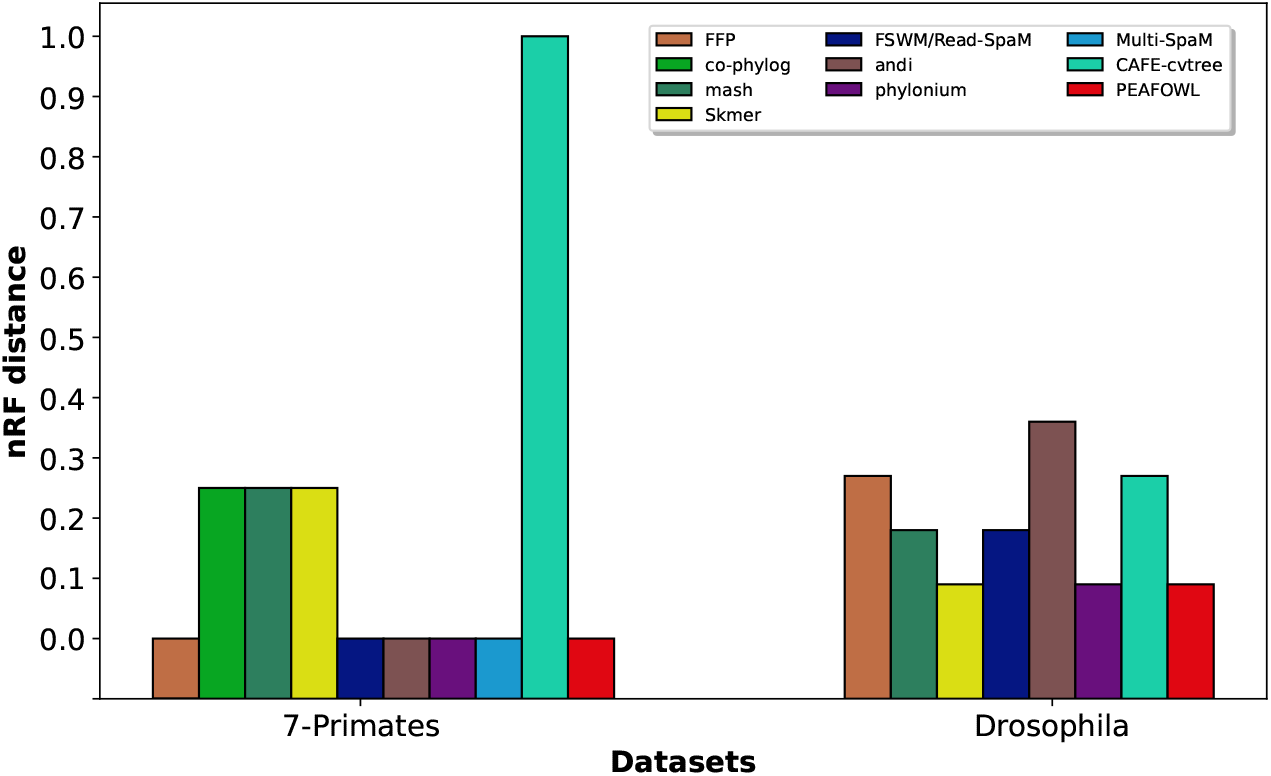
Comparison of nRF distances. nRF distance comparison among Peafowl and state-of-the-art methods on the 7 Primates and Drosophila datasets.

**Figure 5:**
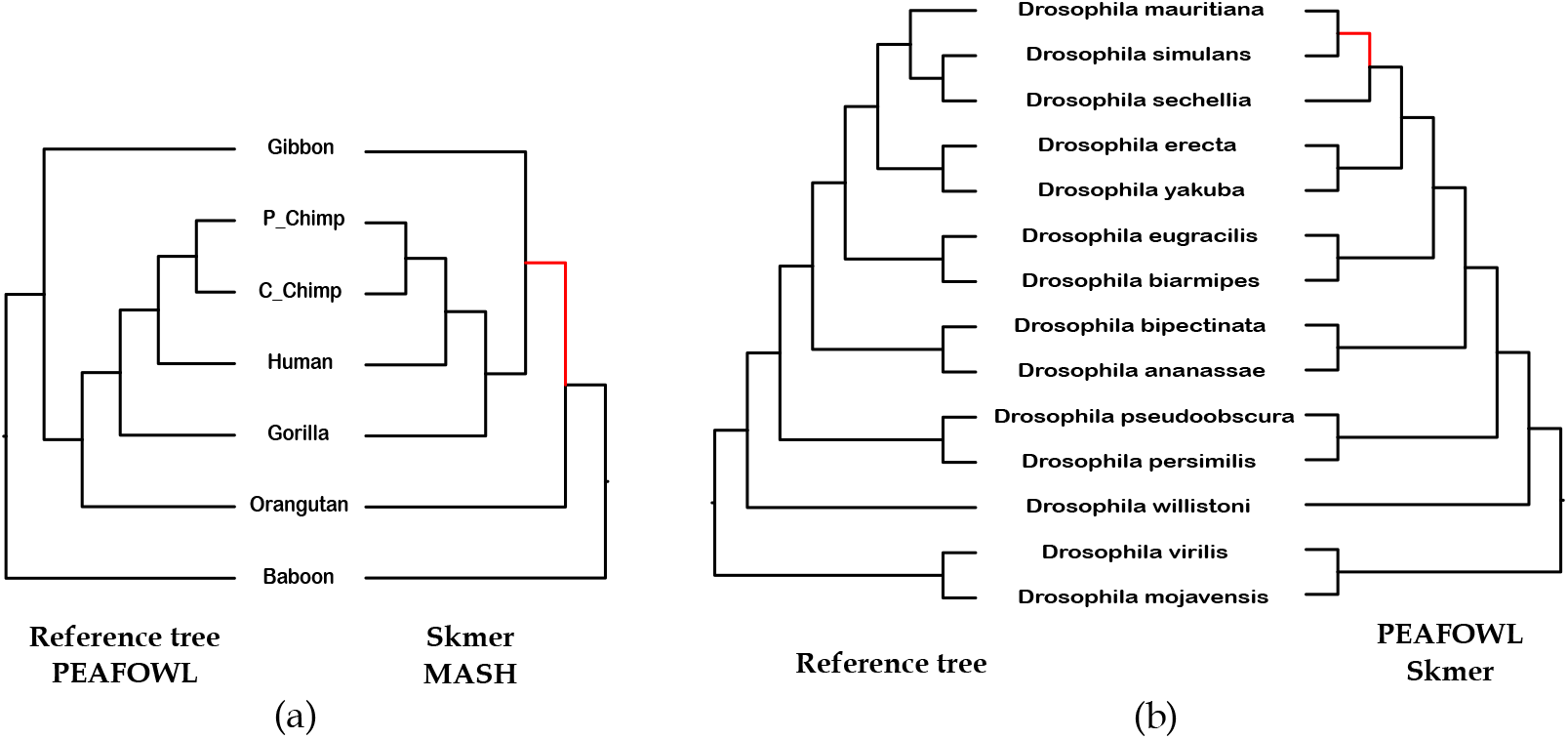
Analysis of the Primate and Drosophila phylogenies. The internal branches in the estimated trees that are not found in the reference trees are shown in red. (a) The trees estimated by Peafowl (which is identical to the reference tree [8]), Skmer, and Mash. (b) The trees estimated by Peafowl and Skmer in comparison to the reference tree.

For the Drosophila dataset, the trees with the lowest nRF distances were obtained by Peafowl, *Skmer* and *phylonium* (Figure 4). Both Peafowl and Skmer produced the same tree which differs from the reference tree in one branch (see Fig. 5(b)). Skmer and Peafowl reconstructed the sister relationship of *Drosophila mauritiana* and *Drosophila simulans* which contradicts the reference tree supporting the (*Drosophila mauritiana*, (*Drosophila simulans, Drosophila sechellia*)) relationship.

### 3.4. Genome-based phylogeny

The Genome-based Phylogeny group of the AFproject [9] includes assembled 29 *E*.*coli/Shigella* strains [10], assembled mitochondrial genomes of 25 fish species of the suborder *Labroidei* [26], and full genome sequences of 14 plant species [27]. A comparison of the nRF distances achieved by various methods is shown in Figure 6. Peafowl attains nRF values of 0.23, 0.05, and 0.36 on these datasets, respectively.

**Figure 6:**
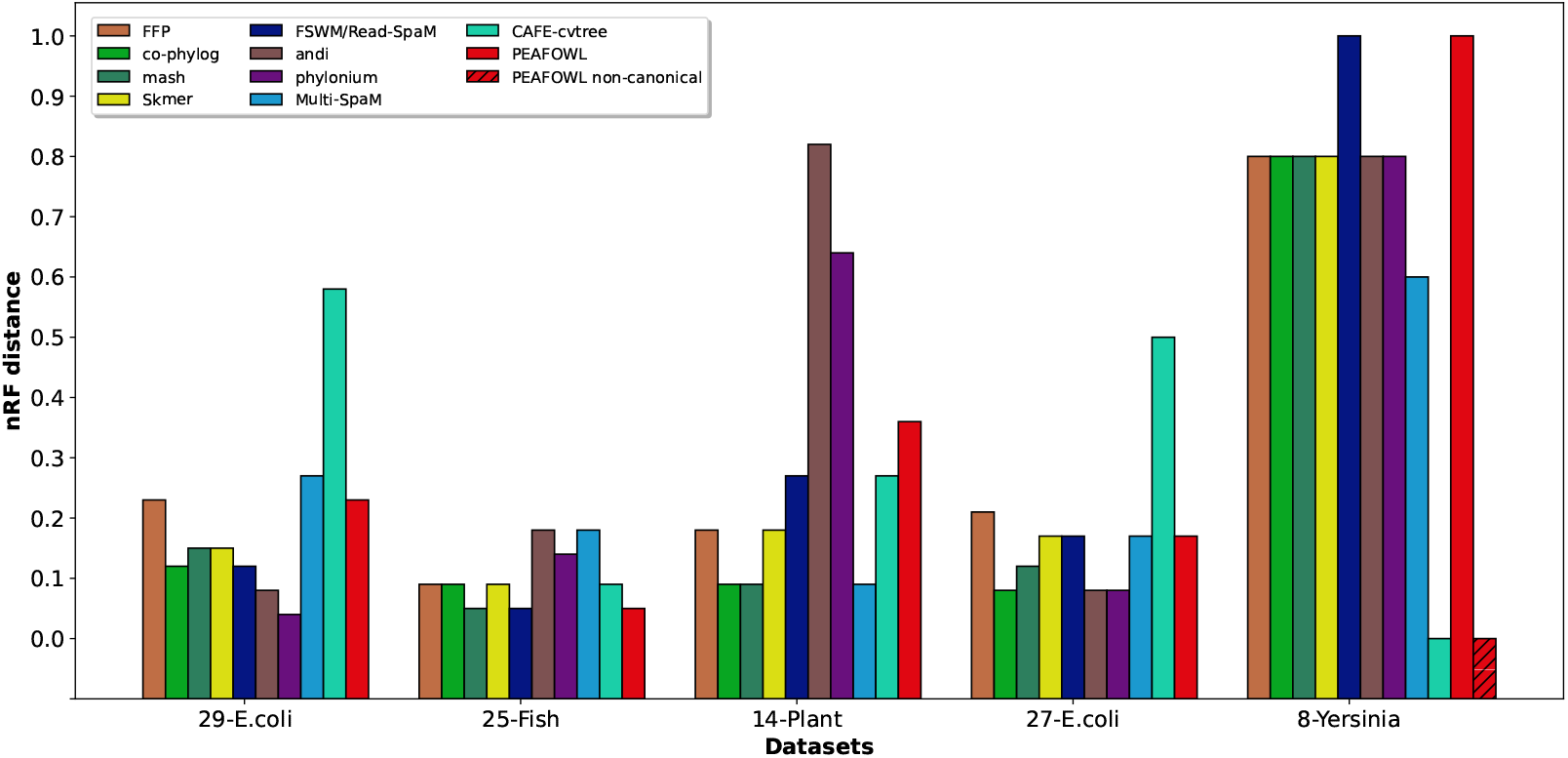
Comparison of nRF distances on the AFproject datasets. nRF distance comparison among Peafowl and several different methods on real datasets from AFproject. Exact values can be found in supplementary materials.

On the 25 Fish dataset, our method is one of the best performing tools, achieving the lowest nRF distance of 0.05 along with *mash* and *FSWM*. Estimated and reference trees for fish genome are shown in S6 Figure.

However, on the 29 *E*.*coli/Shigella* and the 14 plant datasets, Peafowl is outperformed by other methods. The best performing method on the 29 *E*.*coli/Shigella* dataset is phylonium whereas *co-phylog, mash* and *Multi-SpaM* generate the most accurate trees on the 14 plant dataset.

Estimated and reference trees for the 29 *E*.*coli* and 14 plant datasets are available in the supplementary material (S7 Figure, S11 Figure). It is worth noting that the reference tree for the 29 *E*.*coli* dataset was constructed using an alignment-based approach from the assembled genomes [10] and has not been thoroughly validated subsequently. For the plant dataset, the *k*_*entropy*_ value in Peafowl was calculated based on running the method over a *k*-mer range of 9 to 17 instead of 9 to 31 to avoid resource exhaustion. Results are not included in the entropy variation plot for *k* equals 9 and 17 due to the presence of all 1’s in the binary matrix, resulting in zero entropy and computational limitation, respectively.

### 3.5. Horizontal gene transfer (HGT)

This category of data from the AFproject [9] includes full genome sequences of 27 *E*.*coli/Shigella* strains [28] and 8 Yersinia strains [28]. These two datasets are known to have undergone extensive genome rearrangements [9]. They exhibit horizontal gene transfer properties that may cause distant species to show sibling-like properties (such as similar *k*-mers).

The performance of various alignment-free tools on the Yersinia dataset is shown in Figure 6. An observation was made previously [9] that whole-genome analysis tools tend to construct trees relatively discordant to the reference tree on Yersinia sequences than traditional approaches. This seems true for Peafowl as well, with an nRF of 1 on this dataset. The best performing method in this dataset is *CAFE-cvtree*. However, most tools perform poorly in this case, with only two having an nRF value below 0.8. It has been conjectured that the complex nature of the genus and substantial rearrangement events may promote this discrepancy [9].

We further explored this issue and noted that the eight Yersinia genomes are very similar in sequence but share genome rearrangements, and the reference tree was constructed using genomic inversion events inferred from a whole-genome alignment [28]. Since in the canonical counting mode, *k*-mers and their reverse complements are counted together, inversion events are undetected except at the two ends of the inversions. So, we also run our method without the *-r* parameter i.e. perform non-canonical counting, implying that *k*-mers and their reverse complements are treated as separate entities by the counting tool. Remarkably, with the non-canonical counting mode, Peafowl reconstructed a tree that is identical to the reference tree (Fig. 7).

**Figure 7:**
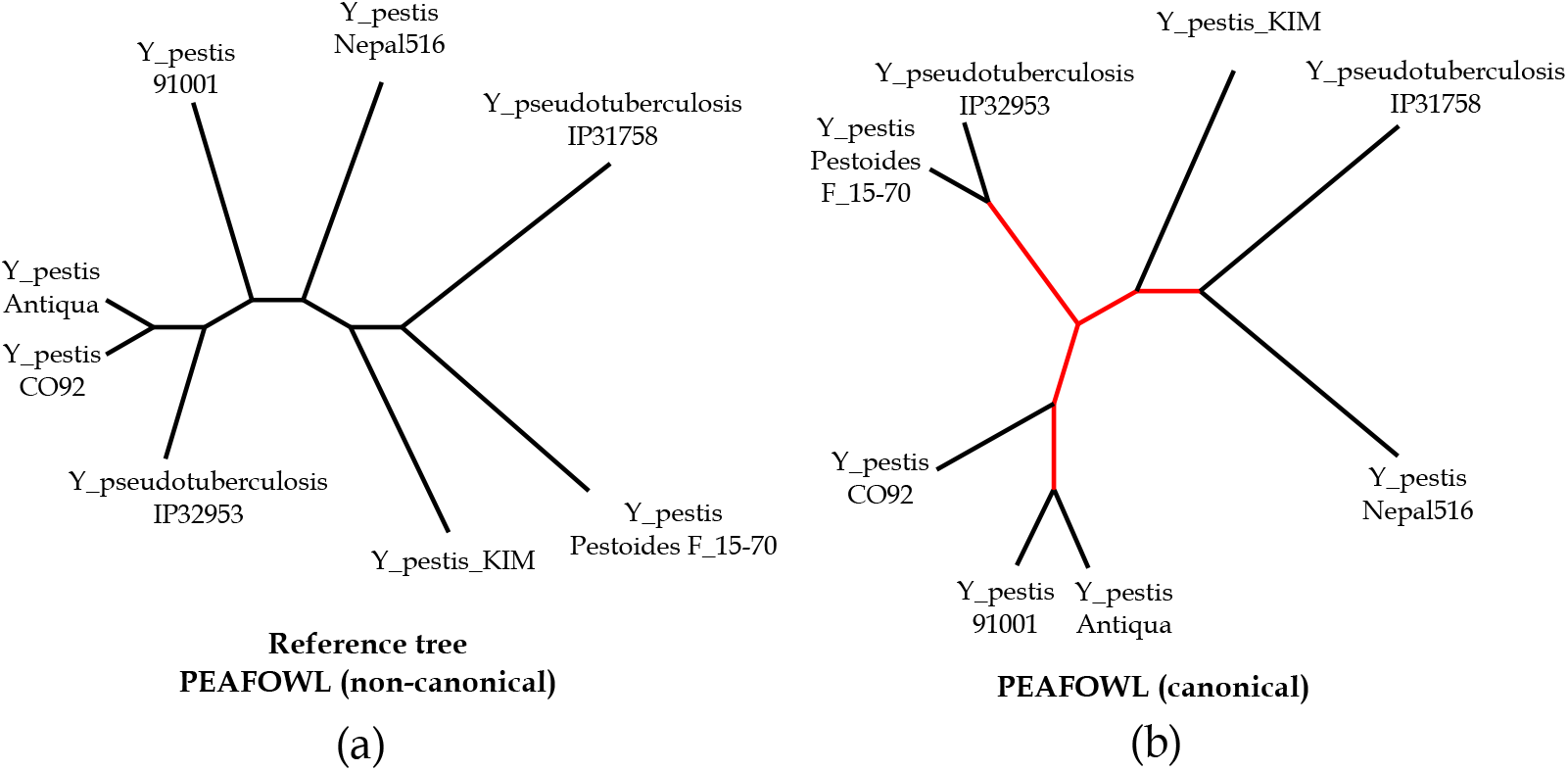
Analysis of the Yersinia phylogenies. (a) The tree estimated by PEAFOWL with non-canonical counting mode, which is identical to the reference tree. (b) PEAFOWL-estimated tree with canonical mode of counting. The branches in the estimated tree that differ from the reference tree are shown in red.

Supplementary S2 Table and S3 Table report the entropy and nRF values (corresponding to the highest entropy) achieved by Peafowl in the canonical and non-canonical settings, respectively. We observe that the entropy values in the non-canonical mode are substantially higher than those in the canonical mode for the Yersinia dataset.

On the 27 *E*.*coli/Shigella* dataset, our method achieves nRF distance of 0.17, but the best performers include *co-phylog, andi* and *phylonium* with an nRF of 0.08 (Figure 6). Estimated and reference trees for *E*.*coli/Shigella* are shown in the supplementary material (S8 Figure).

### 3.6. Runtime and Memory Usage

All the datasets are run in an Intel(R) Core(TM) i7-7700 CPU @ 3.60GHz machine with 8 processors and 32 GB RAM. Table 1 summarizes the unzipped size of the datasets used, corresponding runtime, and peak memory usage by Peafowl. Values for both Plant and Drosophila data are corresponding to a *k*-mer range of 9 to 17.

**Table 1:**
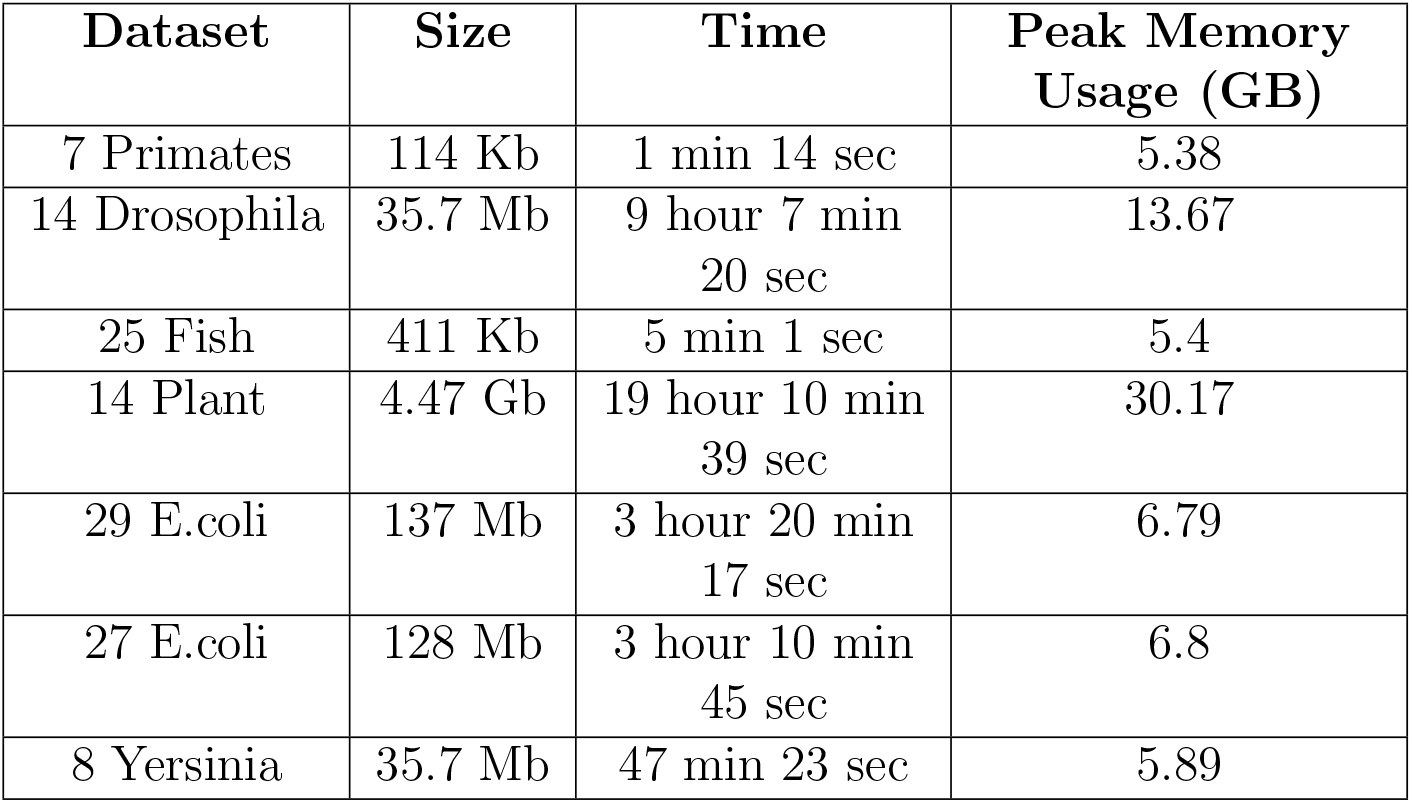
Runtime and peak memory usage of Peafowl.

## 4. Conclusions

In this paper, we presented Peafowl, an alignment-free method for phylogeny estimation using maximum likelihood. It circumvents the complexity of multiple sequence alignment and combines the merits of maximum likelihood estimation in tree construction. We evaluated the performance of Peafowl on seven real datasets and compared the results with the state of the art alignment-free methods. We observe that Peafowl generates trees with the lowest nRF distances in three of the datasets, while phylonium acheives the lowest nRF in four datasets (Figures 4 and 6). Moreover, the tree estimated by Peafowl on one other dataset (Yersinia dataset) matches the reference tree when it is run in a mode suited to capture inversions. Our experimental results suggest that the performance of various methods may substantially vary across different datasets. Therefore, selecting suitable methods becomes particularly challenging when the data are heterogeneous, which is often so for genome-scale phylogenetic data. Consequently, alignment-free tree estimation, far from being a “solved problem”, merits further attention and improvements.

Peafowl has several limitations and can be extended and improved in a number of ways. First, it does not work well if the species are distant since very few *k*-mers are conserved across the species in this case. Its performance also suffers if sequences contain considerable missing regions. In the future, these issues may be addressed. Second, the current version works on assembled sequences or genomes. A future direction will be to extend it to support phylogeny estimation from unassembled sequencing reads. Third, trees are presently estimated based on the presence or absence of *k*-mers, and actual counts are ignored. It may be interesting to explore models to utilize the *k*-mer counts. Finally, our tool is currently designed to handle up to 30 taxa. This limit can be expanded in future to accommodate larger taxonomic groups.

## Supporting information

Supplementary information: an alignment-free method for phylogeny estimation using maximum likelihood

## Appendix A.

### Supplementary Information

Contains following supplementary tables and figures:

*S1 Table*. **Comparison of normalized Robinson Foulds distances**. Comparison of normalized Robinson Foulds distance achieved by different alignment-free methods on real datasets. Minimum nRF values are in bold. For the 7 Primates column, values on the left and right side of oblique represent result obtained using Neighbour joining and UPGMA respectively. For the Yersinia dataset, result of *PEAFOWL* is written as 1/0 denoting nRF of 1 with -r parameter and 0 without.

*S2 Table*. **Canonical entropy values and normalized RF distances obtained by P****eafowl**. Canonical entropy values and normalized RF distances obtained by Peafowl on different datasets. Maximum entropy values are highlighted in bold. Entropy values for 14 plant dataset are reported up to *k*-mer 17 to avoid resource exhaustion.

*S3 Table*. **Non-canonical entropy values and normalized RF distances obtained by P****eafowl**. Non canonical entropy values and normalized RF distances obtained by Peafowl on different datasets. Maximum entropy values are highlighted in bold. Entropy values for 14 plant dataset are reported up to *k*-mer 17 to avoid resource exhaustion.

*S1 Figure*. **Normalized Robinson Foulds distance and entropy vs**. *k***-mer length for the 25-Fish dataset**. Variation of normalized Robinson Foulds distance and entropy with change in *k*-mer length for the 25-Fish dataset. Diamond shaped markers represent values corresponding to *k*_*entropy*_ (*k* = 9).

*S2 Figure*. **Normalized Robinson Foulds distance and entropy vs**. *k***-mer length for the 29 *E*.*coli* dataset**. Variation of normalized Robinson Foulds distance and entropy with change in *k*-mer length for the 29 *E*.*coli* dataset. Diamond shaped markers represent values corresponding to *k*_*entropy*_ (*k* = 23).

*S3 Figure*. **Normalized Robinson Foulds distance and entropy vs**. *k***-mer length for the 14 plant dataset**. Variation of normalized Robinson Foulds distance and entropy with change in *k*-mer length (*k* =11 to 15) for the 14 plant dataset. Diamond shaped markers represent values corresponding to *k*_*entropy*_ (*k* = 15).

*S4 Figure*. **Normalized Robinson Foulds distance and entropy vs**. *k***-mer length for the 27 *E*.*coli* dataset**. Variation of normalized Robinson Foulds distance and entropy with change in *k*-mer length for the 27 *E*.*coli* dataset. Diamond shaped markers represent values corresponding to *k*_*entropy*_ (*k* = 25).

*S5 Figure*. **Normalized Robinson Foulds distance and entropy vs**. *k***-mer length for the 8-Yersinia dataset**. Variation of normalized Robinson Foulds distance and entropy with change in *k*-mer length for the 8-Yersinia dataset (with r parameter). Diamond shaped markers represent values corresponding to *k*_*entropy*_ (*k* = 31).

*S6 Figure*. **Comparison of fish phylogenies. a**. Phylogeny generated by *PEAFOWL* and **b**. the benchmark tree on the Fish dataset.

*S7 Figure*. **Comparison of *E*.*coli* phylogenies. a**. Phylogeny generated by *PEAFOWL* and **b**. the benchmark tree on the 29 *E*.*coli* dataset.

*S8 Figure*. **Comparison of *E*.*coli* phylogenies. a**. Phylogeny generated by *PEAFOWL* and **b**. the benchmark tree on the 27 *E*.*coli* dataset.

*S9 Figure*. **Comparison of Yersinia phylogenies. a**. Phylogeny generated by *PEAFOWL* and **b**. the benchmark tree on the 8 Yersinia dataset.

*S10 Figure*. **Comparison of Yersinia phylogenies (without r parameter). a**. Phylogeny generated by *PEAFOWL* (without r parameter) and **b**. the benchmark tree on the 8 Yersinia dataset.

*S11 Figure*. **Comparison of plant phylogenies. a**. Phylogeny generated by *PEAFOWL* and **b**. the benchmark tree on the 14 Plant dataset.

