## Supplementary information: an alignment-free method for phylogeny estimation using maximum likelihood for "An Alignment-free Method for Phylogeny Estimation using Maximum Likelihood"

| Methods | Datasets | | | | | | |
|---|---|---|---|---|---|---|---|
| | **29 E.coli** | **25 Fish** | **14 Plant** | **27 E.coil** | **8 Yersinia** | **7 Primates** | **Drosophila** |
| FFP | 0.23 | 0.09 | 0.18 | 0.21 | 0.8 | **0** | 0.27 |
| co-phylog | 0.12 | 0.09 | **0.09** | **0.08** | 0.8 | 0.5 | - |
| mash | 0.15 | **0.05** | **0.09** | 0.12 | 0.8 | 0.25/**0** | 0.18 |
| Skmer | 0.15 | 0.09 | 0.18 | 0.17 | 0.8 | 0.25/0.25 | **0.09** |
| FSWM/Read-SpaM | 0.12 | **0.05** | 0.27 | 0.17 | 1 | **0/0** | 0.18 |
| PEAFOWL | 0.23 | **0.05** | 0.36 | 0.17 | 1/**0** | **0** | **0.09** |
| andi | 0.08 | 0.18 | 0.82 | **0.08** | 0.8 | **0/0** | 0.36 |
| phylonium | **0.04** | 0.14 | 0.64 | **0.08** | 0.8 | **0/0** | **0.09** |
| Multi-SpaM | 0.27 | 0.18 | **0.09** | 0.17 | 0.6 | **0** | - |
| CAFE-cvtree | 0.58 | 0.09 | 0.27 | 0.5 | **0** | 0.75/0.25 | - |
| Median AF Project | 0.54 | 0.09 | 0.64 | 0.5 | 0 | - | - |
| Best AF Project | 0.04 | 0.05 | 0.09 | 0.08 | 0 | - | - |

Table S1: **Comparison of normalized Robinson Foulds distances.** Comparison of normalized Robinson Foulds distance achieved by different alignment-free methods on real datasets. Minimum nRF values are in bold. For the 7 Primates column, values on the left and right side of oblique represent result obtained using Neighbour joining and UPGMA respectively. For the Yersinia dataset, result of *PEAFOWL* is written as 1/0 denoting nRF of 1 with -r parameter and 0 without.

| kmer size | Datasets | | | | | | |
|---|---|---|---|---|---|---|---|
| | **7 Primates** | **Drosophila** | **25 Fish** | **29 E.coli** | **14 Plant** | **8 Yersinia** | **27 E.coil** |
| 9 | **3615.96** | 0 | **2573.04** | 79.7426 | 0 | 3.80495 | 77.3329 |
| 11 | 3392.6 | 0.742465 | 1698.89 | 1523.02 | 35.9579 | 279.236 | 1564.18 |
| 13 | 3278.95 | 2146.87 | 1520.86 | 2355.5 | 2340.57 | 698.685 | 2360.82 |
| 15 | 3251.11 | **3241.13** | 1465.35 | 2338.97 | **3410.24** | 842.232 | 2338.36 |
| 17 | 3224.27 | 2370.88 | 1409.93 | 2338.28 | 2456.5 | 846.749 | 2340.6 |
| 19 | 3179.1 | 2172.6 | 1383.88 | 2337.9 | - | 883.142 | 2370.37 |
| 21 | 3157.08 | 2101.94 | 1357.58 | 2310.44 | - | 963.98 | 2354.09 |
| 23 | 3145.58 | 2077.11 | 1330.91 | **2378.96** | - | 1030.61 | 2386.66 |
| 25 | 3107.85 | 2072.25 | 1317.63 | 2362.13 | - | 1040.54 | **2419.14** |
| 27 | 3093.87 | 2056.64 | 1321.92 | 2335.58 | - | 1094.62 | 2396.31 |
| 29 | 3092.59 | 2041.32 | 1303.63 | 2340.9 | - | 1054.41 | 2400.09 |
| 31 | 3067.9 | 2030.61 | 1305.73 | 2351.19 | - | **1117.6** | 2369.13 |
| nRF | 0 | 0.09 | 0.05 | 0.23 | 0.36 | 1 | 0.17 |

Table S2: **Canonical entropy values and normalized RF distances obtained by Peafowl.** Canonical entropy values and normalized RF distances obtained by PEAFOWL on different datasets. Maximum entropy values are highlighted in bold. Entropy values for 14 plant dataset are reported up to $k$-mer 17 to avoid resource exhaustion.

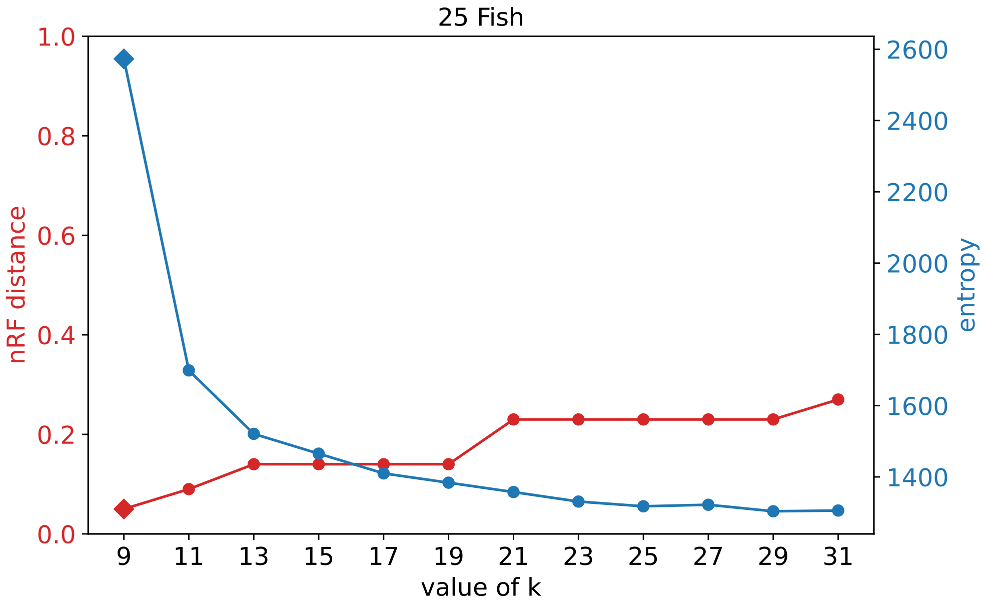


Figure S1: **Normalized Robinson Foulds distance and entropy vs. $k$-mer length for the 25-Fish dataset.** Variation of normalized Robinson Foulds distance and entropy with change in $k$-mer length for the 25-Fish dataset. Diamond shaped markers represent values corresponding to $k_{entropy}$ ($k = 9$).

| kmer size | Datasets | | | | | | |
|---|---|---|---|---|---|---|---|
| | **7 Primates** | **Drosophila** | **25 Fish** | **29 E.coli** | **14 Plant** | **8 Yersinia** | **27 E.coil** |
| 9 | **3575.37** | 0 | **2315.76** | 213.813 | 0 | 87.2463 | 207.321 |
| 11 | 3378.81 | 17.0652 | 1671.15 | 2560.25 | 141.889 | 2955.31 | 2574.37 |
| 13 | 3292.69 | 3244.44 | 1517.93 | **2640.69** | 3115.87 | 4400.54 | **2645.71** |
| 15 | 3259.07 | **3469.01** | 1449.79 | 2433.96 | **3165.45** | 4498.11 | 2473.8 |
| 17 | 3210.92 | 2210.08 | 1406.96 | 2370.21 | 2259.79 | **4502.69** | 2393.63 |
| 19 | 3181.13 | 2064.89 | 1377.99 | 2355.63 | - | 4491.7 | 2387.07 |
| 21 | 3165.69 | 2013.31 | 1346.19 | 2328.45 | - | 4502.33 | 2386.53 |
| 23 | 3137.06 | 2002.93 | 1331.23 | 2314.3 | - | 4496.69 | 2364.63 |
| 25 | 3116.49 | 1991.83 | 1332.8 | 2350.04 | - | 4459.67 | 2346.08 |
| 27 | 3091.58 | 1967.92 | 1320.05 | 2319.48 | - | 4463.77 | 2341.95 |
| 29 | 3086.13 | 1985.55 | 1303.31 | 2282.54 | - | 4467.69 | 2339.97 |
| 31 | 3071.44 | 1970.84 | 1300.85 | 2295.84 | - | 4431.96 | 2290.89 |
| nRF | 0 | 0.18 | 0.05 | 0.62 | 0.46 | 0 | 0.58 |

Table S3: **Non-canonical entropy values and normalized RF distances obtained by Peafowl.**Non canonical entropy values and normalized RF distances obtained by PEAFOWL on different datasets. Maximum entropy values are highlighted in bold. Entropy values for 14 plant dataset are reported up to $k$-mer 17 to avoid resource exhaustion.

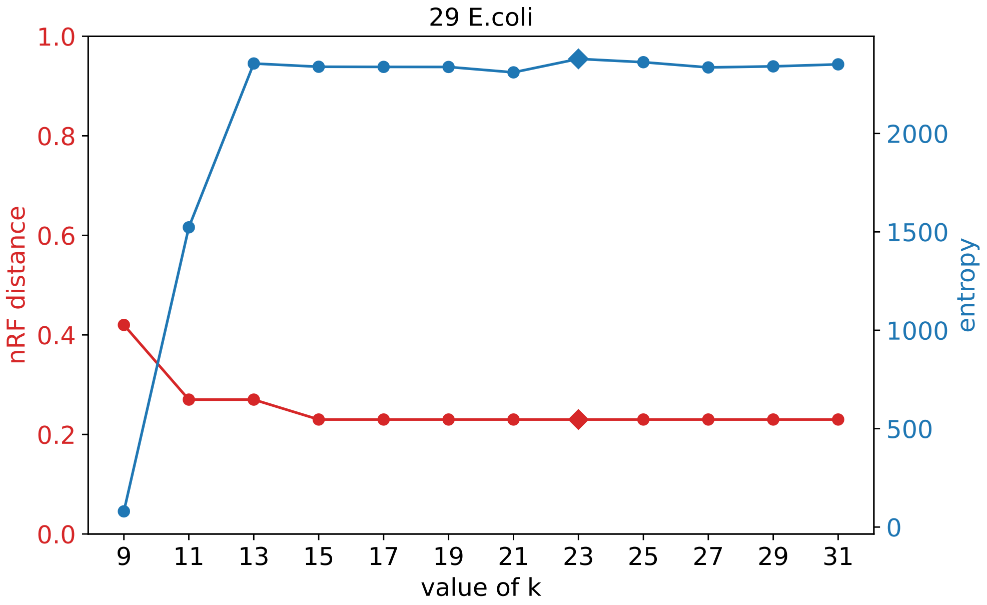


Figure S2: **Normalized Robinson Foulds distance and entropy vs. $k$-mer length for the 29 *E.coli* dataset.** Variation of normalized Robinson Foulds distance and entropy with change in $k$-mer length for the 29 *E.coli* dataset. Diamond shaped markers represent values corresponding to $k_{entropy}$ ($k = 23$).

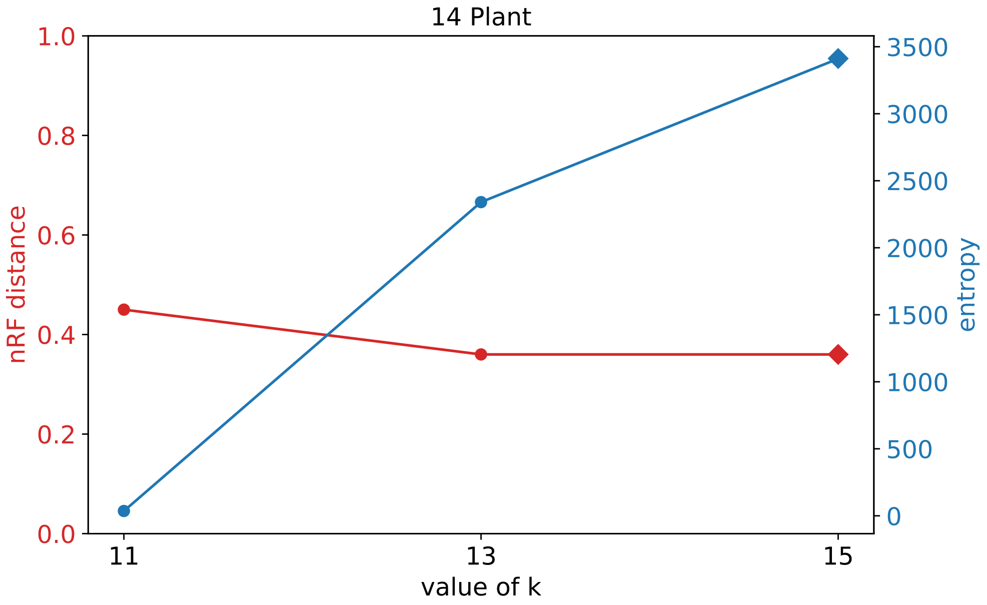


Figure S3: **Normalized Robinson Foulds distance and entropy vs. $k$-mer length for the 14 plant dataset.** Variation of normalized Robinson Foulds distance and entropy with change in $k$-mer length ($k$=11 to 15) for the 14 plant dataset. Diamond shaped markers represent values corresponding to $k_{entropy}$ ($k$ = 15).

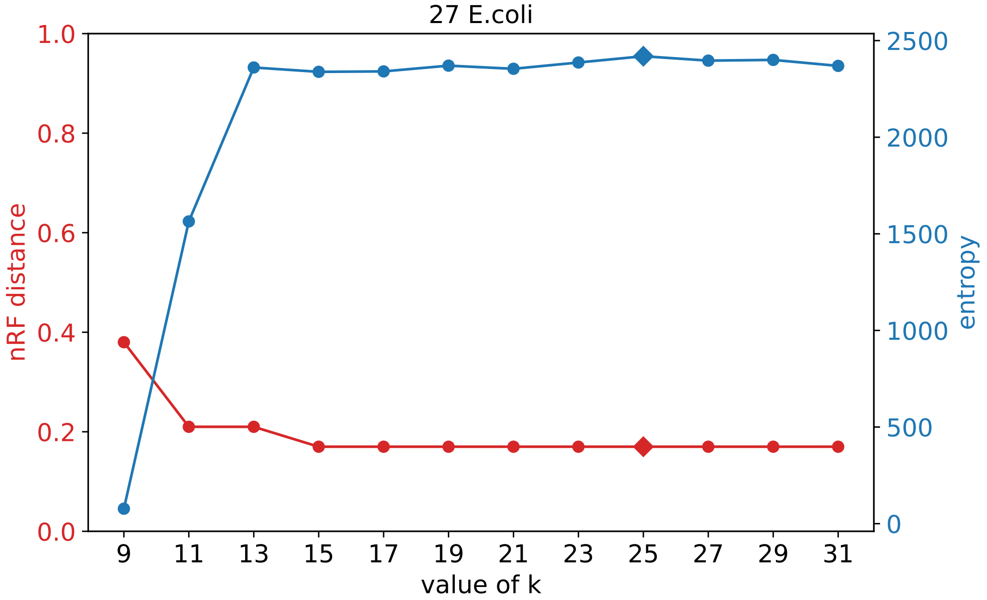


Figure S4: **Normalized Robinson Foulds distance and entropy vs. $k$-mer length for the 27 *E.coli* dataset.** Variation of normalized Robinson Foulds distance and entropy with change in $k$-mer length for the 27 *E.coli* dataset. Diamond shaped markers represent values corresponding to $k_{entropy}$ ($k$ = 25).

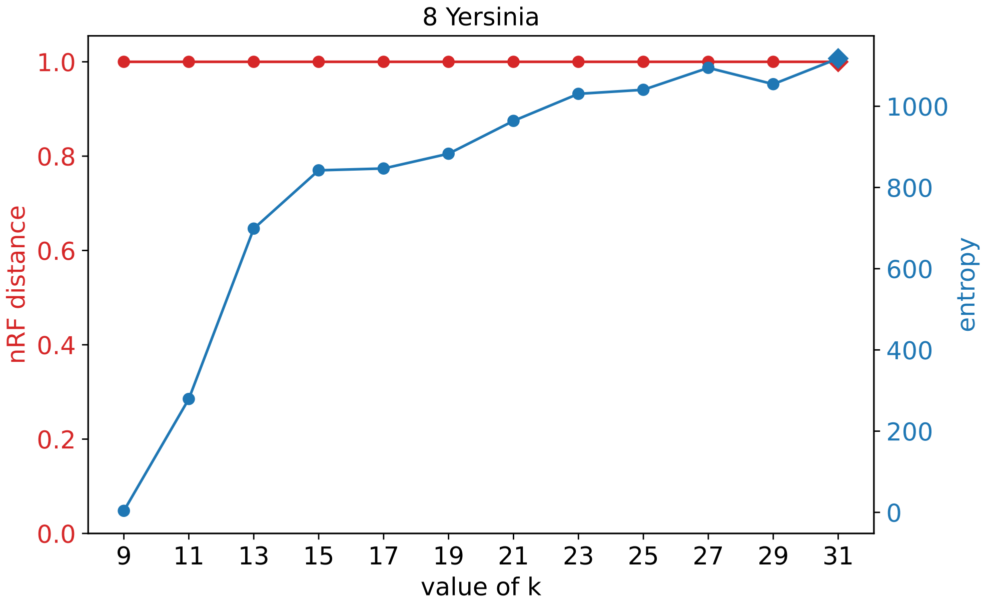


Figure S5: **Normalized Robinson Foulds distance and entropy vs. $k$-mer length for the 27 8-Yersinia dataset.** Variation of normalized Robinson Foulds distance and entropy with change in $k$-mer length for the 8-Yersinia dataset (with r parameter). Diamond shaped markers represent values corresponding to $k_{entropy}$ ($k = 31$).

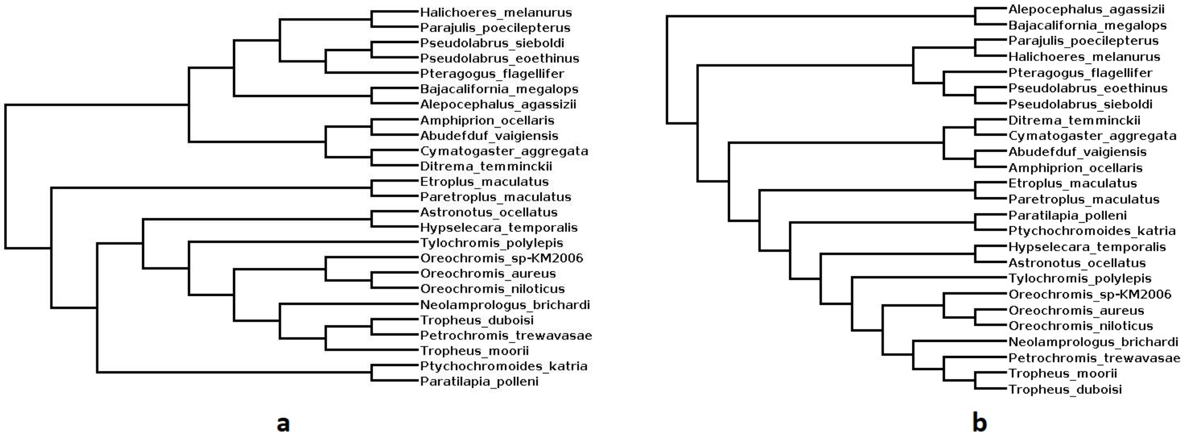


Figure S6: **Comparison of fish phylogenies. a.** Phylogeny generated by *PEAFOWL* and **b.** the benchmark tree on the Fish dataset.

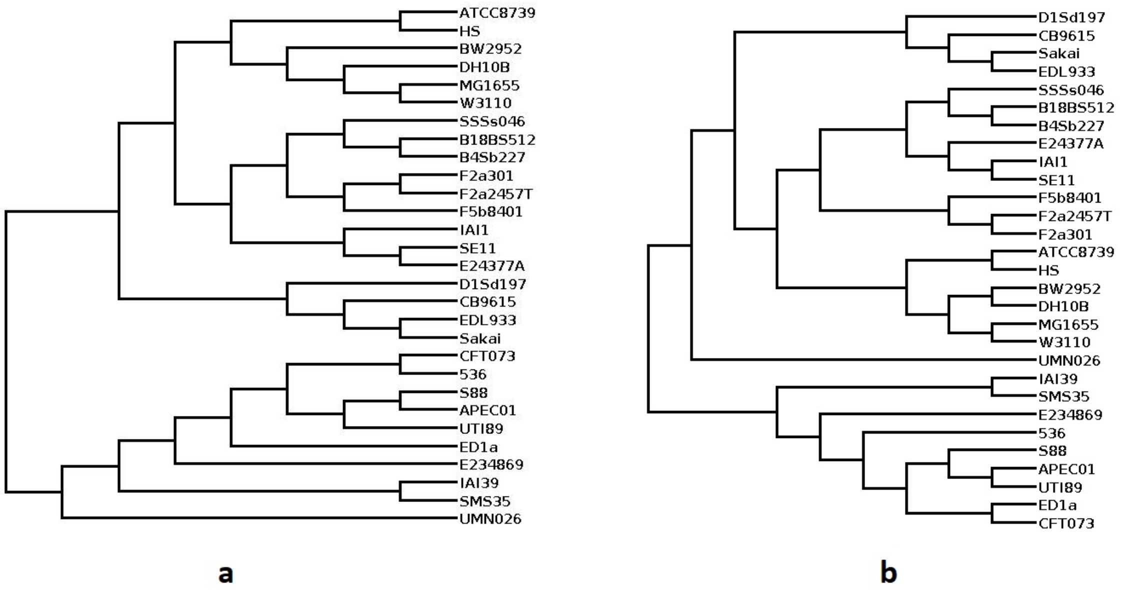


Figure S7: **Comparison of *E.coli* phylogenies. a.** Phylogeny generated by *PEAFOWL* and **b.** the benchmark tree on the 29 *E.coli* dataset.

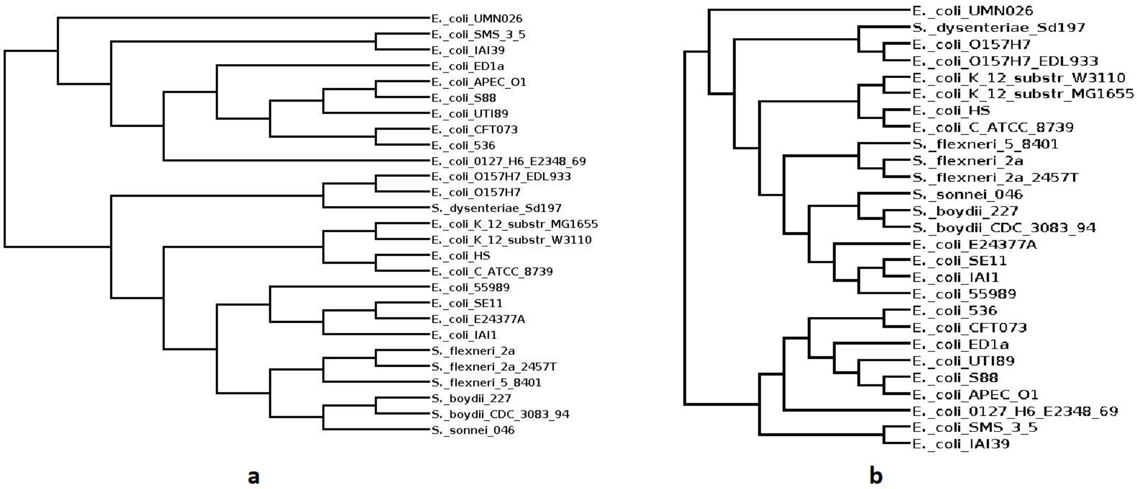


Figure S8: **Comparison of *E.coli* phylogenies. a.** Phylogeny generated by *PEAFOWL* and **b.** the benchmark tree on the 27 *E.coli* dataset.

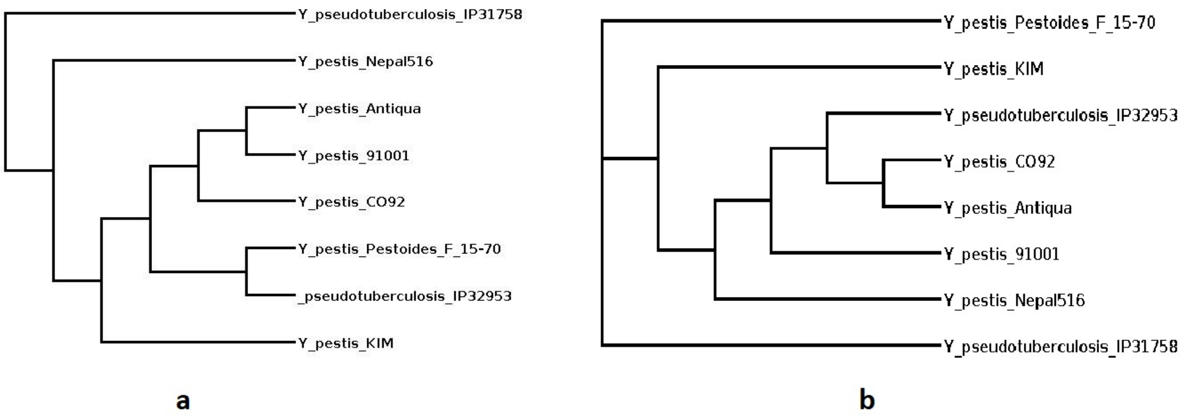


Figure S9: **Comparison of Yersinia phylogenies. a.** Phylogeny generated by *PEAFOWL* and **b.** the benchmark tree on the 8 Yersinia dataset.

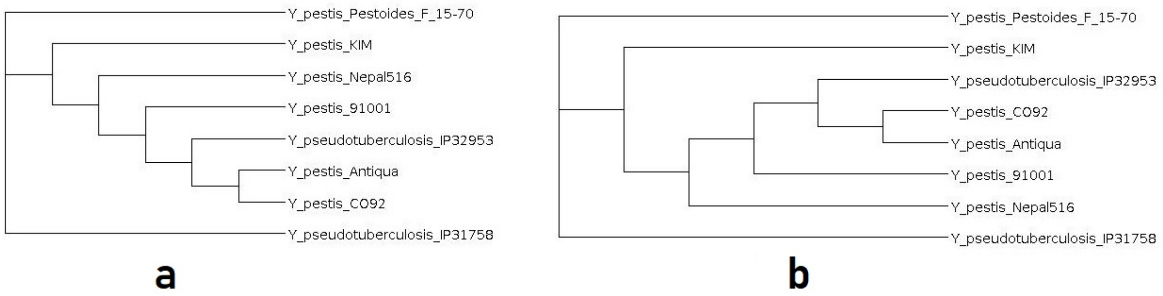


Figure S10: **Comparison of Yersinia phylogenies (without r parameter). a.** Phylogeny generated by *PEAFOWL* (without r parameter) and **b.** the benchmark tree on the 8 Yersinia dataset.

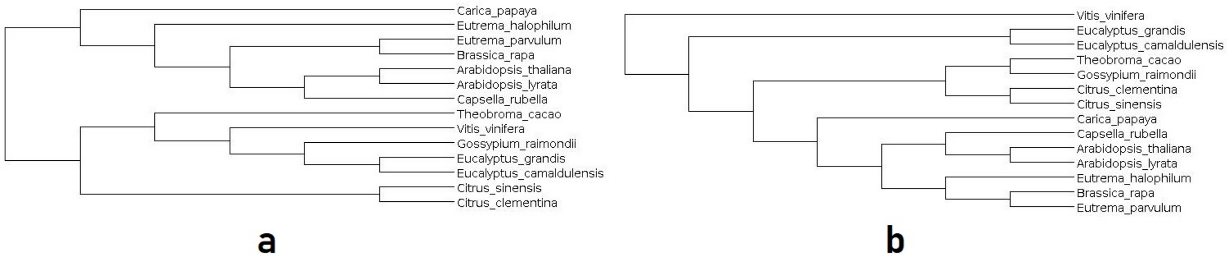


Figure S11: **Comparison of plant phylogenies. a.** Phylogeny generated by *PEAFOWL* and **b.** the benchmark tree on the 14 Plant dataset.
